# Synaptopodin is required for long-term depression at Schaffer collateral-CA1 synapses

**DOI:** 10.1101/2023.08.21.554176

**Authors:** Yanis Inglebert, Pei You Wu, Julia Tourbina-Kolomiets, Cong Loc Dang, R. Anne McKinney

## Abstract

Synaptopodin (SP), an actin-associated protein found in telencephalic neurons, affects synaptic and structural plasticity. While being required for longterm depression (LTD) mediated by metabotropic glutamate receptor (mGluR-LTD), little is known about its role in other forms of LTD induced by low frequency stimulation (LFS-LTD) or spike-timing dependent plasticity (STDP). Using SP-deficient mice (SPKO), we have here demonstrated that the presence of SP is mandatory for normal LTD expression. On the contrary, long-term potentiation (LTP), albeit diminished in SPKO, is still present and can be easily recovered by increasing the stimulation frequency. Our study shows, for the first time, the role of SP in a more physiological form of induction of synaptic plasticity.

## Introduction

Synaptopodin (SP) is a postsynaptic actin-associated protein found in a subset of mature dendritic spines ^1^. Mainly found in the hippocampus and cortex, it is known to be involved in synaptic plasticity and spine remodeling ^2,3^. For example, lack of SP has been shown to result in a reduced long-term potentiation (LTP) and deficits in spatial learning ^4^. More recently, our group has demonstrated that SP is required for a form of long-term depression (LTD) mediated by group I metabotropic glutamate receptor (mGluR-LTD) ^5,6^. However, it is unclear whether SP is also required for other forms of LTD. Additionally, two distinct forms of plasticity induction based on synaptic activity coexist: Bienenstock-Cooper-Munroe (BCM) model and Spike-Timing Dependent Plasticity.

The first is a theoretical framework describing how synaptic plasticity can be modulated by the level of neuronal activity and the history of synaptic inputs ^7^. In this paradigm, low-frequency stimulation (2 – 5 Hz) induces LTD (LFS-LTD) while high-frequency stimulation (10 – 100 Hz) induces LTP (HFS-LTP) but the threshold between LTD/LTP can be dynamically adjusted. Both require an increase in intracellular calcium (Ca^2+^) from NMDA receptors (NMDAR), at least at CA3-CA1 synapses ^8^. The role of SP in this form is still poorly understood. While some studies suggested that only spines containing SP are more likely to develop LFS-LTD ^9^, others reported a normal LFS-LTD in SP-deficient mice ^10^.

The second model is called Spike-Timing Dependent Plasticity (STDP). More physiological, rather than being dependent on the stimulation frequency, synaptic modifications are based on the precise timing between presynaptic (in the form of excitatory postsynaptic potential or EPSP) and postsynaptic (in the form of an action potential or AP) activity. These paradigms are thought to better mimic what happens in the brain ^11^. Classically, timing-dependent long-term potentiation (t-LTP) is induced when an EPSP is followed by an AP in the postsynaptic neuron ^12,13^. In contrast, timing-dependent long-term depression (t-LTD) is induced when an EPSP is preceded by an AP ^14–16^. At CA3-CA1 synapses, both induction protocols also require intracellular calcium elevation, but while t-LTP requires strong Ca^2+^ entry from NMDA- and AMPA-receptors (AMPAR) activation, t-LTD relies on moderate Ca^2+^ release from internal stores through IP3 receptors (IP3R) triggered by mGluR5 activation ^17,18^. This is a general learning rule underlying memory found *in vitro* and *in vivo* in a wide range of species ^19^ but to date no study has explored the role of SP in STDP.

In this study, to better understand the role of SP at Schaffer collateral-CA1 synapses (Sc-CA1), we induced LFS-LTD/HFS-LTP or t-LTD/t-LTP in wild-type (WT) and SP-deficient mice (SPKO). Our results show that in the absence of SP, it is impossible to induce LFS-LTD or t-LTD. Nevertheless, although affected, t-LTP can be recovered by boosting Ca^2+^ entry through specific activity patterns. These results provide novel insight into the role of SP in LTD and describe SP as a key molecular component to enable synaptic plasticity.

## Results

### Synaptopodin is required for LTD induced by low frequency stimulation

First, we wanted to determine the consequences of the absence of SP on synaptic plasticity induced by low-or high-frequency stimulation to gain an overview of the BCM curve. The Schaffer collaterals were stimulated with 900 pulses at 2, 10 or 100 Hz (**Fig 1A**). LFS-LTD and HFS-LTP are known to be induced respectively by 2 and 100 Hz while 10 Hz is generally the threshold between LTD/LTP. In SPKO, no significant change in synaptic modification (field EPSP slope measured on the last 10 minutes of the recording) was observed after 2 Hz (100.3 ± 1,7 vs 81.4 ± 6.1, p < 0.003, Mann-Whitney) and 10 Hz (110.68 ± 7.8 vs 152.96 ± 11.1, p < 0.03, Mann-Whitney) stimulation compared to WT (**Fig 1C, D**). In contrast, WT exhibited a robust LFS-LTD and HFS-LTP respectively for 2 Hz and 10 Hz stimulation. At 100 Hz, SPKO revealed a modest HFS-LTP in comparison to the magnitude found in WT (**Fig 1E**; 117.9 ± 3.4 vs 140.1 ± 3.7, p < 0.001, Mann-Whitney). Overall, in SPKO, our data suggest that LFS-LTD is absent and HFS-LTP amplitude is decreased (**Fig 1B**). Interestingly, the threshold for synaptic changes appears to be higher in the absence of SP.

**Figure 1.**
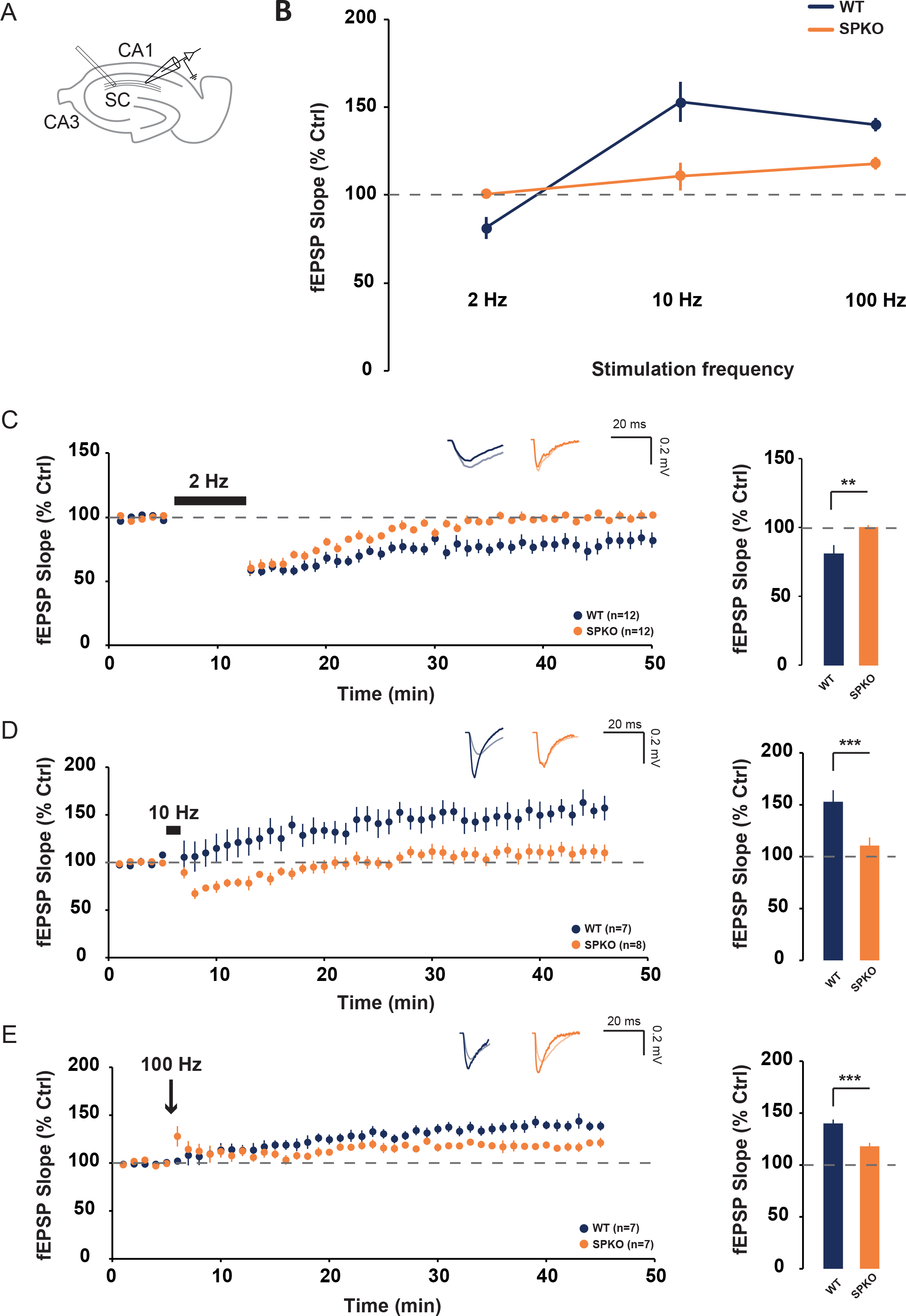
SPKO mice show a deficit of LFS-LTD and HFS-LTP. **(A)** Diagram showing the positions of stimulating and recording electrodes at the Sc-CA1 synapses. **(B)** BCM curves in WT and SPKO for 2, 10 and 100 Hz stimulation frequency. **(C)** *(Left)* Time course of normalized fEPSP in WT and SPKO following 2Hz (900 pulses) stimulation. *(Right)* Quantification of average synaptic modification in the last 10 minutes of the recording. **(D)** *(Left)* Time course of normalized fEPSP in WT and SPKO following 10Hz (900 pulses) stimulation. *(Right)* Quantification of average synaptic modification in the last 10 minutes of the recording. **(E)** *(Left)* Time course of normalized fEPSP in WT and SPKO following 100Hz (900 pulses) stimulation. *(Right)* Quantification of average synaptic modification in the last 10 minutes of the recording.

### Spike-timing dependent plasticity is altered in absence of synaptopodin

We next examined the consequences of the lack of SP on synaptic plasticity induced at the Schaffer collateral-CA1 pyramidal cell synapses by STDP (**Fig 2A, Left**). Classically, t-LTP was induced by repeatedly pairing (100 times, 0.3 Hz) an EPSP followed briefly (between +5 and +50ms, positive pairing) by a postsynaptic action potential (AP) (**Fig 2A, Right**). No t-LTP is observed in SPKO at +10ms, a timing that generally shows the highest t-LTP amplitude in WT. (**Fig 2B**, 83.3 ± 8,6 vs 139.2 ± 7,1, p < 0.001, Mann-Whitney). Instead, a robust t-LTD is observed. Construction of STDP curves for several timings exhibited a single LTD window ranging from +10ms to +30ms in SPKO (**Fig 2C, Right**) while the conventional LTP window is observed in WT (**Fig 2C, Left**). For negative pairing (a postsynaptic AP followed by an EPSP) at -25ms, a timing that generally shows the highest t-LTD amplitude in WT, no plasticity is observed in SPKO (**Fig 2D**, 96.2 ± 3.7 vs 71.9 ± 7.7, p < 0,05, Mann-Whitney). Rather, no plasticity is observed. These data indicate that the STDP rule is severely impaired in the absence of SP and t-LTD is totally absent for negative pairings.

**Figure 2.**
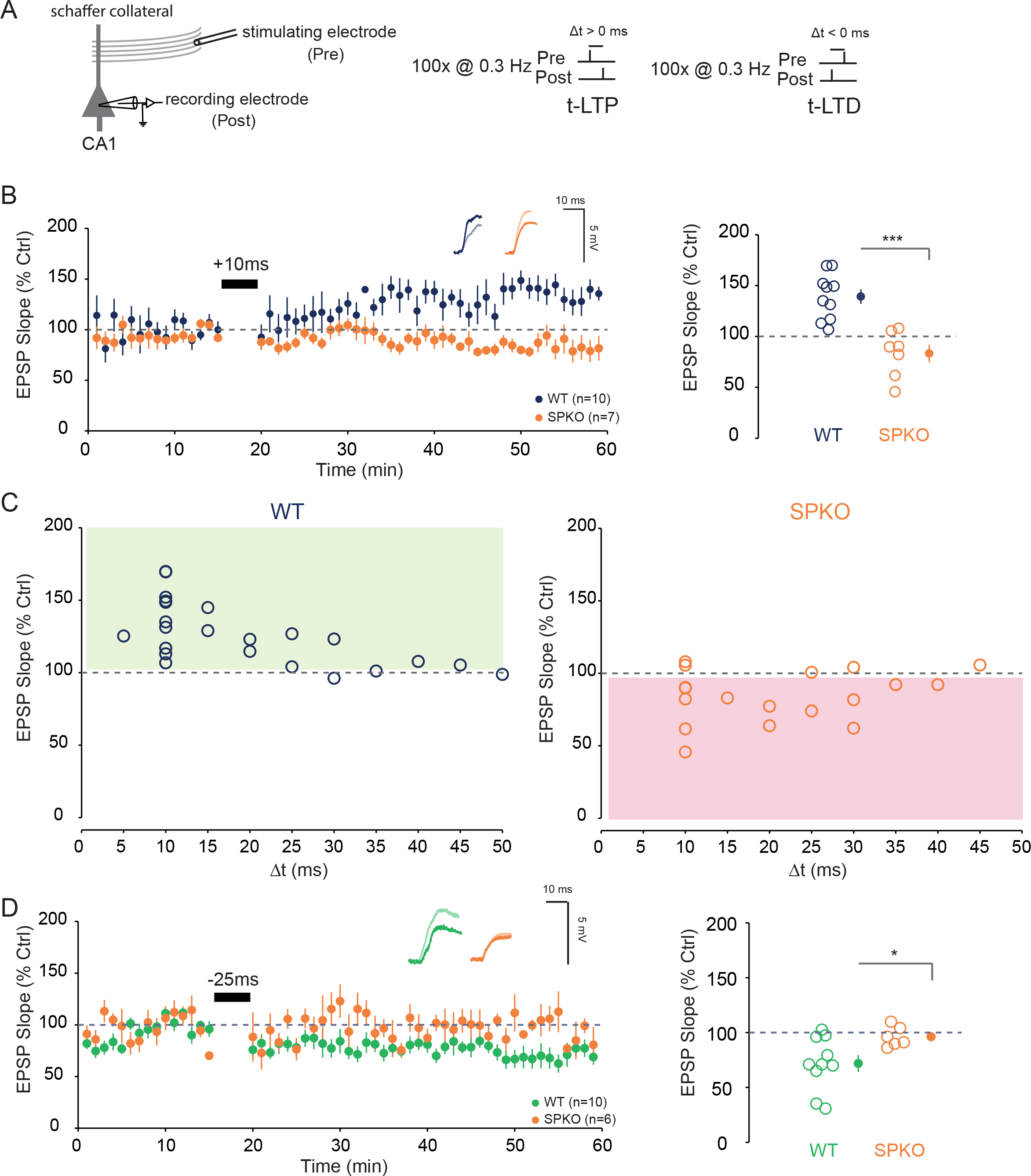
Spike-Timing Dependent Plasticity is altered in SPKO. **(A)** *(Left)* Diagram showing the positions of stimulating electrodes and recording electrodes at the Sc-CA1 synapses. *(Right)* Illustration of the pre-before-post (100p@0.3Hz) and post-before-pre (100p@0.3Hz) protocol used to induce synaptic modification. **(B)**. *(Left)* Time course of normalized EPSP in WT and SPKO following pre-beforepost protocol at +10ms. *(Right)*. Quantification of average synaptic modification (*closed circles*) in the last 10 minutes of the recording with individuals’ values for each cell (*open circles*). **(C)**. STDP curves in WT and SPKO for pre-before-post protocol at different timings. **(D)**. *(Left)* Time course of normalized EPSP in WT and SPKO following post-before-pre protocol at -25ms. *(Right)*. Quantification of average synaptic modification (*closed circles*) in the last 10 minutes of the recording with individuals’ values for each cell (*open circles*).

### t-LTP but not t-LTD can be rescued in absence of synaptopodin

STDP induction rules are flexible and can be adapted by modulating intracellular Ca^2+^ influx ^20–22^. For example, an easy way to boost intracellular Ca^2+^ influx is to increase the stimulation frequency which would lead to cross the synaptic threshold for t-LTD and t-LTP. Taking this into account we attempted to recover t-LTP and t-LTD deficits in SPKO by increasing the pairing frequency. At +10ms, increasing the pairing frequency from 0.3 to 5 Hz has been sufficient to significantly recover the t-LTP window (**Fig 3A**, 129.9 ± 10.3, p < 0,05, MannWhitney). On the other hand, increasing the pairing frequency did not significantly recover t-LTD window at -25ms neither at 5 Hz (101.6 ± 5.7) nor at 10 Hz (87.7 ± 3.7) compared to 0.3 Hz (**Fig 3B**). Similarly to the results obtained in **Fig1**, t-LTP, while affected, can still be induced but t-LTD remained totally absent. We concluded that t-LTP can be rescued in SPKO but not t-LTD.

**Figure 3.**
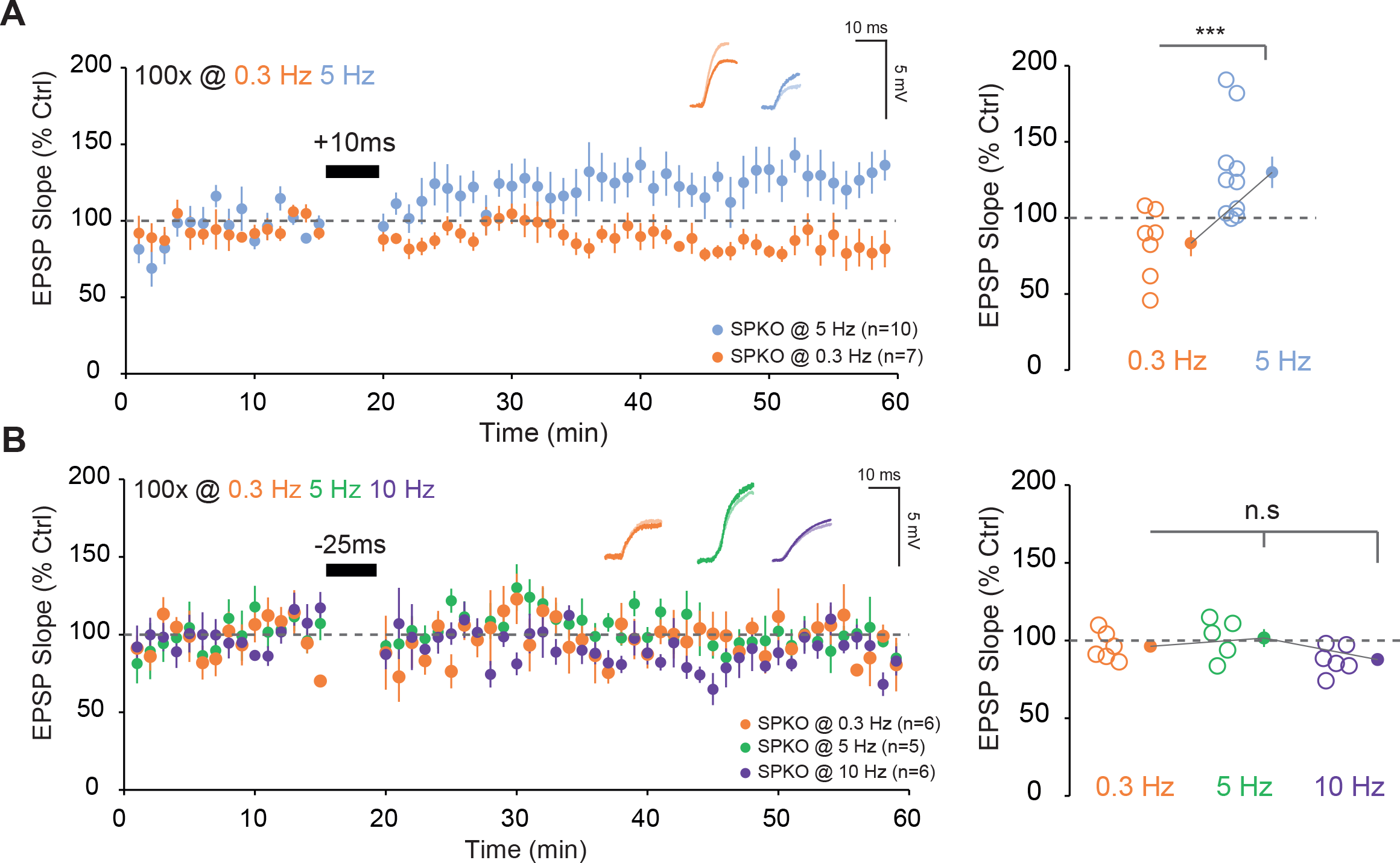
t-LTP but not t-LTD can be restored in SPKO. **(A)**. *(Left)* Time course of normalized EPSP in WT and SPKO following pre-before-post (+10ms) protocol at 0.3 or 5 Hz. *(Right)*. Quantification of average synaptic modification (*closed circles*) in the last 10 minutes of the recording with individuals’ values for each cell (*open circles*). **(B)**. *(Left)* Time course of normalized EPSP in WT and SPKO following post-before-pre (-25ms) protocol at 0.3, 5 or 10 Hz. *(Right)*. Quantification of average synaptic modification (*closed circles*) in the last 10 minutes of the recording with individuals’ values for each cell (*open circles*).

### AMPA and IP3-receptors expression is altered in SPKO

To better understand what could lead to these deficits of synaptic plasticity, we next examined the expression level of the key receptors involved in these two forms of synaptic plasticity. NMDA, AMPA and IP3-receptors are the main sources of calcium to induce synaptic changes. Their expression level was analyzed by Western blot from hippocampi lysates of WT and SPKO. We particularly investigated GluA1/GluA2 subunit of AMPARs and GluN2A/GluN2B subunit of NMDARs because they are the major subunits found in the hippocampus ^23,24^. Significant decrease was observed for total expression of AMPA-receptor subunits GluA1 (1.07±0.04 vs 0.81±0.03, p<0.01, unpaired t-test) and GluA2 (1.05±0.03 vs 0.92±0.02, p=0.01, unpaired t-test) in SPKO compared to WT (**Fig 4A**). Similar results were found regarding the surface expression of GluA1 (**Fig 4B**, 0.50±0.04 vs 0.37±0.04, p<0.05, unpaired t-test) and GluA2 (**Fig 4C**, 0.46±0.04 vs 0.33±0.03, p<0.01, unpaired t-test). In contrast, no difference in expression was found for NMDA receptor subunits GluN2A (**Fig 4D Left**, 0.90±0.03 vs 0.89±0.04, p>0.05, unpaired t-test) and GluN2B (**Fig 4D Right**, 0.94±0.04 vs 1.01±0.01, p>0.05, unpaired t-test). Finally, IP3-receptors expression was found to be significantly decreased in SPKO (**Fig 4E**, 1.00±0.06 vs 0.78±0.05, p<0.05, unpaired t-test). Overall, IP3- and AMPA-receptors expression was found to be significantly reduced unlike NMDA receptors.

**Figure 4.**
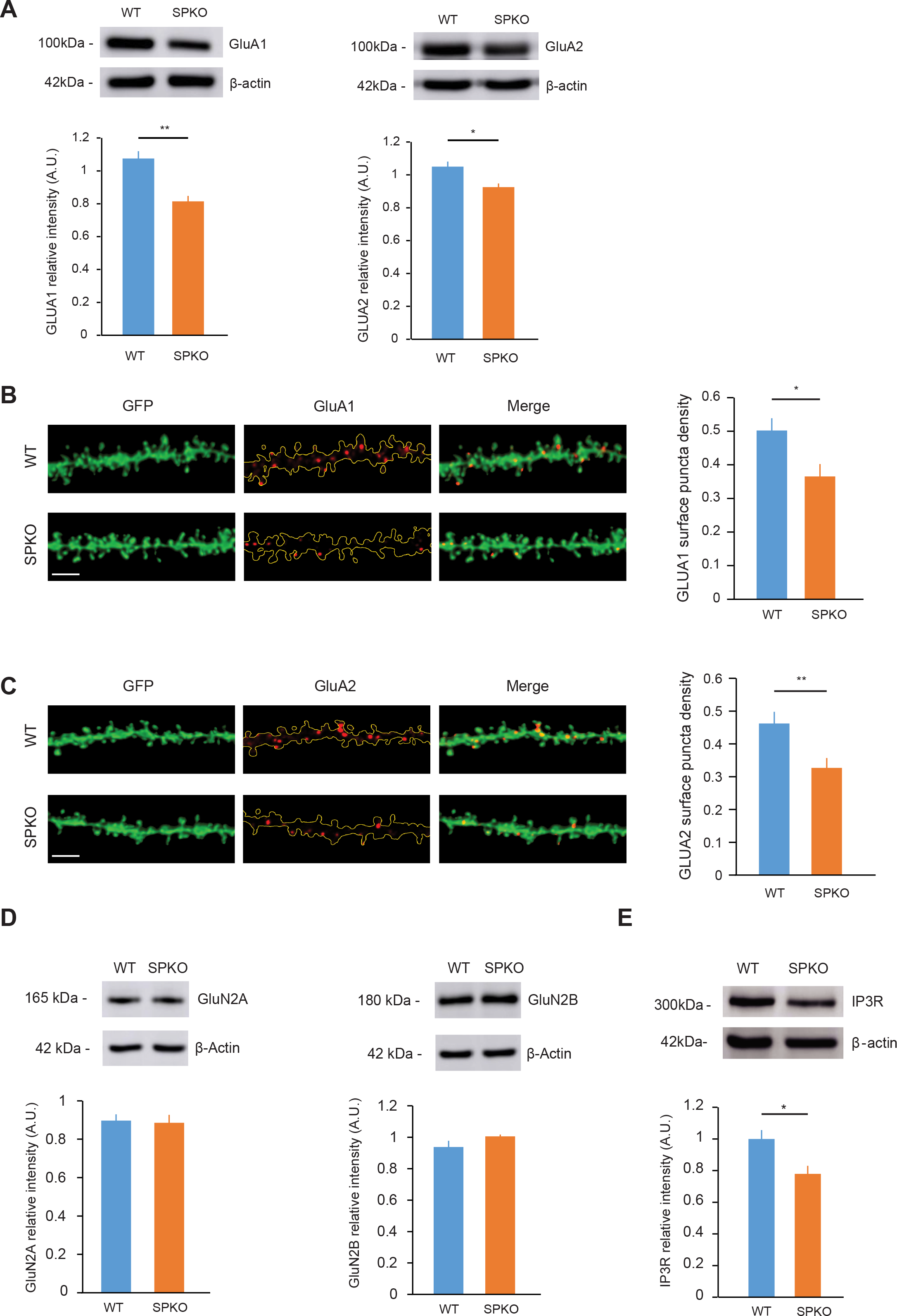
GluA1, GluA2 and IP3R are decreased in SPKO. **(A)**. Representative immunoblot and quantification of total (*left*) GluA1 and (*right*) GluA2 from hippocampi of WT (n=5) and SPKO (n=5) mice. **(B) (C)**. Representative images and quantification of CA1 pyramidal neuron tertiary dendrites immunostained with surface GluA1 antibody (WT n=12, KO n=14) and surface GluA2 antibody (WT n=13, KO n=11) in acute hippocampal slices. Scale bar=3 μm. **(D)**. Representative immunoblot and quantification of total (left) NR2A and (right) NR2B from hippocampi of WT (n=5) and SPKO (n=5) mice. **(E)**. Representative immunoblot and quantification of total IP3R from hippocampi of WT (n=8) and SPKO (n=8) mice.

## Discussion

### LFS-LTD is absent in SPKO

Our results revealed an absence of LFS-LTD in SPKO compared to WT (**Fig 1**). Our findings contradict previously published findings ^10^ that LFS-LTD was not modulated by loss of SP. The experimental conditions are similar regarding the stimulation protocol (2Hz 1200 stimuli vs 2Hz 900 stimuli) and the age of mice (P14-P25). An attractive explanation for this difference could be that the LFS-LTD threshold is higher in SPKO. In fact, previous studies have shown a relationship between the number of stimuli and LTD amplitude ^25^. Another possible explanation is that the extracellular calcium concentrations used are different. While we used a more physiological concentration of calcium (2mM), Zhang et al, employed a concentration of 2.5mM. Higher calcium concentrations could lead to overestimation of plasticity ^21,26^.

### BCM curve is shifted in SPKO

In SPKO slices, our data showed that at 10 Hz, HFS-LTP was absent. 10 Hz is a particularly interesting frequency, as it is considered to be the threshold (θm) between LTD and LTP ^7^. This threshold is dynamic and slidable and can serve as a homeostatic mechanism to keep synapses within a modifiable zone in response to perturbations in neuronal activity. Interestingly, lack of SP has been linked to a loss of GABAergic inhibition ^27^ and increased excitability ^28^. To compensate for this increase in CA1 cell excitability, the sliding threshold could horizontally shift to the right as demonstrated in other brain regions ^29^. Further studies are required to better understand the role of SP in homeostatic plasticity at CA3-CA1 synapses. One elegant recent paper identified SP as a molecular tag to promote input-specific synaptic plasticity ^30^.

### t-LTP can be rescued but not t-LTD

At Sc-CA1 synapses, both t-LTP and t-LTD require post-synaptic calcium (Ca^2+^) elevation ^31,32^ but from different sources. The magnitude of the Ca^2+^ signal determines the direction/polarity of plasticity ^33^. Whereas t-LTP relies almost solely on Ca^2+^ entry via post-synaptic NMDA receptors (NMDAR), t-LTD required activation of mGluR5 and subsequent IP3R-mediated Ca^2+^ release from internal stores ^17^ and both are affected by the absence of SP. As we previously reported, mGluR5 function and expression is impaired ^6^. Moreover, SP is known to regulate release of Ca^2+^ from internal stores ^34^ and our results indicate that IP3R expression is impaired. In other words, if t-LTD is absent and impossible to recover, it’s certainly because the needed molecular machinery is not present in the absence of SP. In contrast, t-LTP can simply be recovered by increasing the pairing frequency (from 0.3 to 5 Hz) and therefore the post-synaptic calcium entry. In addition to NMDAR, AMPA receptors mediated EPSPs provide local depolarization to boost NMDAR current ^35,36^. A reduction of AMPAR expression could reduced NMDAR current for a given depolarization explaining the absence of t-LTP observed at 0.3 Hz. A stronger protocol (5 Hz) enables the potentiation threshold to be crossed because NMDAR expression is not affected in SPKO, and the calcium entry provided by NMDAR only is now sufficient. Additional calcium imaging experiments will be required to better understand the calcium dynamics at the level of the single spine-expressing SP.

Our results demonstrate for the first time the central role of SP in LTD, particularly induced by STDP. These findings encourage further exploration and dissection of the role of SP in plasticity at the scale of the single spine. It could be the key to understanding why dendritic spines are not all equal when it comes to plasticity.

## Materials and Methods

### Animals

All procedures were according to protocols approved by the guidelines of the Canadian Council on Animal Care and the McGill University Comparative Medicine and Animal Resources animal handling protocols 5057. C57Bl6 (WT) or SPKO mice were used as previously described ^2,22^. Mice were fed *ad libitum* and housed with a 12 h light/dark cycle. WT and SPKO male and female mice were used for experiments.

### Electrophysiology

Hippocampal *ex vivo* slices were obtained from P14-P25 old WT or SPKO mice. Mice were deeply anesthetized with isoflurane and killed by decapitation. Slices (400 μm or 350 μm) were cut on a vibratome (Leica Microsystems, VT1200S) in a sucrose-based solution containing the following (in mM): 280 sucrose, 26 NaHCO3, 10 glucose, 1.3 KCl, 1 CaCl2, and 10 MgCl2 and were transferred at 32°C in regular ACSF containing the following (in mM): 124 NaCl, 5 KCl, 1.25 NaH2PO4, 2 MgSO4, 26 NaHCO3, 2 CaCl2, and 10 glucose saturated with 95% O2/5% CO2 (pH 7.3, 300 mOsm) for 15 min before resting at room temperature (RT) for 1 h in oxygenated (95% O2/5% CO2) ACSF. Recordings were obtained using a Axopatch 200B (Molecular Devices) amplifier and pClamp10.4 software. Data were sampled at 10 kHz, filtered at 3 kHz, and digitized by a Digidata 1440A (Molecular Devices). Field excitatory postsynaptic potentials (fEPSPs) or excitatory postsynaptic potentials (EPSPS) were recorded in the stratum radiatum of the CA1 region by using glass microelectrodes filled with 3 M NaCl. fEPSPs or EPSPs were elicited at 0.1 Hz by a digital stimulator that fed by a stimulation isolator unit. All data analyses were performed with custom-written software in Igor Pro 8 (Wavemetrics). fEPSP or EPSP slope was measured as an index of synaptic strength. For whole-cell recordings, access resistance was monitored throughout the recording and only experiments with stable resistance were kept (changes<20%) and recordings were made from CA1 pyramidal neurons, electrodes were filled with a solution containing the following (in mM): 120 K-gluconate, 20 KCl, 10 HEPES, 2 MgCl26H2O, and 2 Na2ATP.

### Immunohistochemistry

Hippocampal slices of 100 μm were obtained from P30 to P40 old L15S or L15 mice. The slices were incubated in ACSF at RT for 1 h for recovery and fixed in 0.1 M Phosphate buffer (PB) containing 4% PFA, pH 7.4, overnight at 4°C. After fixation, slices were washed in 0.1 M PB and blocked with 1.5% heat-inactivated horse serum overnight at 4°C. Slices were incubated with primary anti-GLUA1 antibody (1:200, Alomone Lab, CAT# AGC-004) and primary anti-GLUA2 antibody (1:200, Alomone Lab, CAT# AGC-005) in blocking solution for 5 days at 4°C, washed with PB, and incubated with anti-rabbit secondary antibody conjugated to DyLight 649 (1:500; Jackson ImmunoResearch Laboratories) overnight. They are then washed and mounted with DAKO Fluorescent Mounting medium (Dako Canada) onto microscope slides before imaging and subsequent blinded analysis. Spine images were taken in z stacks using a Leica Microsystems SP8 confocal microscope with oil-immersion 63x objective at 6x zoom-in. The images were deconvolved and analyzed using software Huygens (Scientific Volume Imaging) and Imaris (Oxford Instruments) respectively. The experimenter was blinded to the treatment.

### Immunoblotting

Hippocampi of WT and SPKO mice at P20 were obtained and rapidly homogenized in ice-cold RIPA (150 mM NaCl, 1% Nonidet P-40, 0.5% sodium deoxycholate, 0.1% sodium dodecylsulfate, 50 mM Tris-HCl (pH 8.0) with protease and phosphatase inhibitors (1X Roche Complete Mini, 5 mM NaF, 1 mM sodium orthovandate, 1 mM PMSF). Homogenates were sonicated and centrifuged for 5 min at 13,200 rpm. The supernatant was extracted and mixed with 2x Laemmli sample buffer to produce loading samples. Samples were boiled, resolved on SDS-PAGE, transferred to nitrocellulose, and stained with primary antibodies overnight. The following primary antibodies were used: GluA1 (1:1000, Abcam, ab31232), GluA2 (1:1000, Abcam, ab20673), GluN2A (1:1800, Abcam, ab124913,), GluN2B (1:1800, Abcam, ab65783), IP3R (1:1000, Cell Signaling, D53A5). Horseradish peroxidase (HRP)-conjugated secondary antibodies goat anti-rabbit (abcam; 1:5000) or goat anti-mouse (Bio-Rad Laboratories; 1:10,000) were applied for 1 h. The blots were imaged with Amersham Imager 600 and the band intensities were analyzed using the open-source program Fiji.

## Acknowledgments and funding sources

Canadian Institutes of Health Research MOP 86724 to R.A.M.; NSERC Discovery RGPIN-2020-06373 to R.A.M. and the Norman Zavalkoff Family Foundation to R.A.M.; Richard and Edith Strauss Postdoctoral Fellowship in Medicine to Y.I.; NSERC-USRA to J.T-K. We thank members of R.A.M. laboratories for comments on the manuscript; and Francois Charron for excellent technical assistance. The authors declare no competing financial interests.

## Author contributions

Y.I. and R.A.M. designed research; Y.I., P.Y.W, C.L.D and J.T-K. performed research; Y.I., P.Y.W., C.L.D. and J.T-K. analyzed data; Y.I., P.Y.W and R.A.M. wrote the paper.

